# A Comparison of Comorbidities and their Contribution to Medical Resource Utilization for Matched HIV-Infected and Uninfected Individuals: A Cross-Sectional Analysis

**DOI:** 10.1101/678136

**Authors:** Michelle Odlum, Sunmoo Yoon

**Affiliations:** Columbia University Medical Center, New York, NY 10032

**Keywords:** Charlson Scores, Comorbidities, Electronic Health Records, Medical Resource Utilization

## Abstract

HIV long-term survivorship is characterized by higher rates of comorbidities compared to uninfected groups. Aging with HIV involves complex interactions of factors (e.g., Individual Characteristics, Infections) that result in a 20% increase in comorbidity risk. With over half of the 1.1 million people living with HIV in the US age 50 and over, the need exists to further understand this interplay and differences in aging-related outcomes. Electronic health record data was analyzed for HIV infected (N=208) and uninfected (N=208) adult inpatients, propensity score matched by age and gender. Diagnostic codes were extracted that comprise the factors of Individual Characteristics, High Risk Behaviors, Chronic Conditions, Mental Health Conditions and Infections. Identified codes were assessed for their contributions to medical resource utilization, based on Charlson Comorbidity scores. Significant contributors to high Charslon scores for HIV infected patients were age (β=0.116; [95% CI 0.077, 0.155]) and admission frequency (β=0.159; [95% CI 0.114, 0.205]) in addition to the comorbidities of acute kidney failure (β=3.27; [95% CI 1.76, 4.78]), hypertension (β= −1.77; [95% CI −2.99, −0.551]). Significant contributors for HIV uninfected patients were age (β=0.110; [95% CI 0.087, 0.133]), length of hospital stay (β=0.006; [95% CI 0.003, 0.009]), acute kidney failure (β=1.556; [95% CI 0.611, 2.50]), heart failure (β= 1.713; [95% CI 0.717, 2.71]), and diabetes mellitus II (β= 1.385; [95% CI 0.634, 2.14]). Our findings enhance the understanding of the contributions to medical resource utilization based on HIV status and can inform intervention efficacy for improved HIV aging outcomes.

## Introduction

There is a growing US population of aging persons living with HIV/AIDS (PLWH) because of diagnosis in later life or long-term survivorship [1,2]. Immune restoration with highly active antiretroviral therapy (HAART) [1,3] has contributed tremendously to these outcomes. However, long-term survivorship is characterized by the presence of and elevated risk for comorbidities [4, 5] not simply explained by the decline in AIDS-related morality and longer life [2,3,6]. Research has linked HIV infection to conditions including cardiovascular disease [7–10]. After adjusting for usual risk factors, HIV-associated rates remain the highest for PLWH who are younger, indicating accelerated aging. In fact, a recent study concluded that the HIV virus has the potential to accelerate aging by more than 14 years [11]. Therefore, aging with HIV must be further explored. The determinants of aging with HIV include complex interactions of comorbid factors including biological/clinical (e.g., diabetes) and socio-behavioral (e.g., smoking). With half of the 1.1 million PLWH, in the US estimated to be 50 or over [7,13], understanding and improving aging of PLWH is a priority.

Aging phenotype development has improved the understanding of physical and cognitive decline in populations of aging adults with multiple comorbidities [16]. Several positive aging phenotypes are characterized (e.g., physical and social functioning) to allow investigators to study avenues for healthier aging outcomes [17]. Although a frailty phenotype is proposed in middle aged HIV infected women, the application of phenotypes to HIV infection is understudied. More effective HIV interventions in aging are dependent on identifying narrower phenotypes with greater clinical validity [17]. However, barriers exist to understanding the interplay between HIV and aging. These are attributed to difficulties in comparing HIV subgroups to the general population of HIV uninfected adults. With this limited understanding of the link between aging in PLWH and uninfected groups [18,19], the current study seeks to fill this gap. Moreover, the recent widespread adoption of electronic health records (EHR) in the US has afforded us the opportunity to leverage clinical data to further HIV phenotype development.

Utilizing EHR data, we investigated the contributions to medical resource utilization based on differences in Charlson comorbidity scores. We sought to understand and classify between group differences in electronic clinical data of HIV infected and uninfected controls, propensity score matched on gender and age. Our cross-sectional study looked at the presence of comorbidities and not HIV-related contributions (e.g., disease stage and immune status) to the development or proliferation of comorbidities. Higher Charlson scores are an indication of the increased likelihood the predicted outcome will result in either 1-year mortality or the higher use of medical resources, [20] the current study focused on the later. In this paper, we report the Individual Characteristics, High Risk Behaviors, Chronic Conditions, Mental Health Conditions and Infections that predict high Charlson scores by HIV status. Findings will enhance our understanding of aging with HIV for effective disease management and improved outcomes in HIV infected populations.

## Materials and Methods

### Patient Population

HIV infected (N=208) and uninfected (N=208) inpatient records were matched based on age and gender for adults 18 and older between January of 2006 and December of 2014, from a clinical data warehouse of electronic health records (EHR). Institutional review board approval was obtained to analyze the de-identified data, which excluded all potentially identifiable patient information (e.g., name, address, date of birth). No patients were involved in the data analysis or interpretation. After data cleaning, we were left with a total of 16,334: HIV infected N=208) and uninfected (N=l 9,216) patients for matching. Mahalanobis’ propensity scoring was used to match HIV infected patients to comparable HIV uninfected patients [17]. Matching allows for meaningful comparisons between two groups and reduces confounding factors in the statistical assessment of outcomes. Diagnostic codes utilized in our analysis were extracted from encounters, past histories, and problem lists. Factors were identified as Individual Characteristics (e.g., ICD 9/10: 262 - Malnutrition), High Risk Behaviors (e.g., ICD9/10: 305.1 – use of Tobacco), Chronic Conditions (e.g., ICD9/10: 584.9 - Acute Kidney Failure), Mental Health Conditions (e.g., ICD9/10: 311- Depressive Disorders), and Infections (e.g., ICD9/10: 070.41- Hepatitis C). Group inclusion was based on the presence or absence of a diagnosis in the chart history.

### Statistical Analysis

The association between Charlson scores and Individual Characteristics, High Risk Behaviors, Chronic and, Mental Health Conditions and Infections were examined. Findings are summarized using descriptive statistics. Pearson Product Moment Correlations (PPMCs) were calculated to determine the relationship between variables that comprise identified factors and Charlson scores (an indicator of medical resource utilization). T-tests were used to assess differences in continuous variables and chi square analyses were used to assess differences in categorical variables. Two independent stepwise multiple regressions (i.e., HIV+ and HIV-) were performed to identify the relevant importance of identified variables (p<0.05) to high Charlson scores. The stepwise approach allows for the prevention of bias in the selection of variables in the final models [21, 22]. Betas (β) and confidence intervals (CIs) were reported for regression analyses, with the use of SPSS 23.0.

## Results

A total of 416 patients were included in our analysis, ages 18 to 85, with the mean age of 50.6±13.2. The racial distribution of the HIV infected sample (N=208) includes 27.1% (N=56) Blacks; 21.4% (N=45) Whites; 0.64% (N=1) Asian; 14.3% (N=30) Mixed Race; 12.46% (N=26) Other; and24.1% (N=50) Unknown or Declined. The HIV uninfected sample (N=208) includes 10.1% (N=21) Blacks; 31.3% (N=65) Whites; 3.1% (N=6) Asian; 15.4% (N=32) Mixed race; 10.8% (N=22) Other; and 29.3% (N=61) Unknown or Declined. The frequency of patient admissions was 7.57±10.7 for HIV patients and 6.17±8.4 for HIV uninfected patients. The average length of stay for HIV patients was 86.6±159.5 and 63.4± 105 for HIV uninfected patients. The top four ICD9/10 codes for the HIV patients were Substance Abuse, Hypertension, Hyperlipidemia and Hepatitis C whereas, the top four for HIV uninfected patients were Hypertension, Diabetes Mellitus II, history of tobacco use and Substance abuse, Table 1. Charlson scores ranged from 0-21 with an average of 7.42±4.35 for HIV patients and 0-12 with an average of 3.52±2.97 for HIV uninfected patients, Table 2.

**Table 1.**
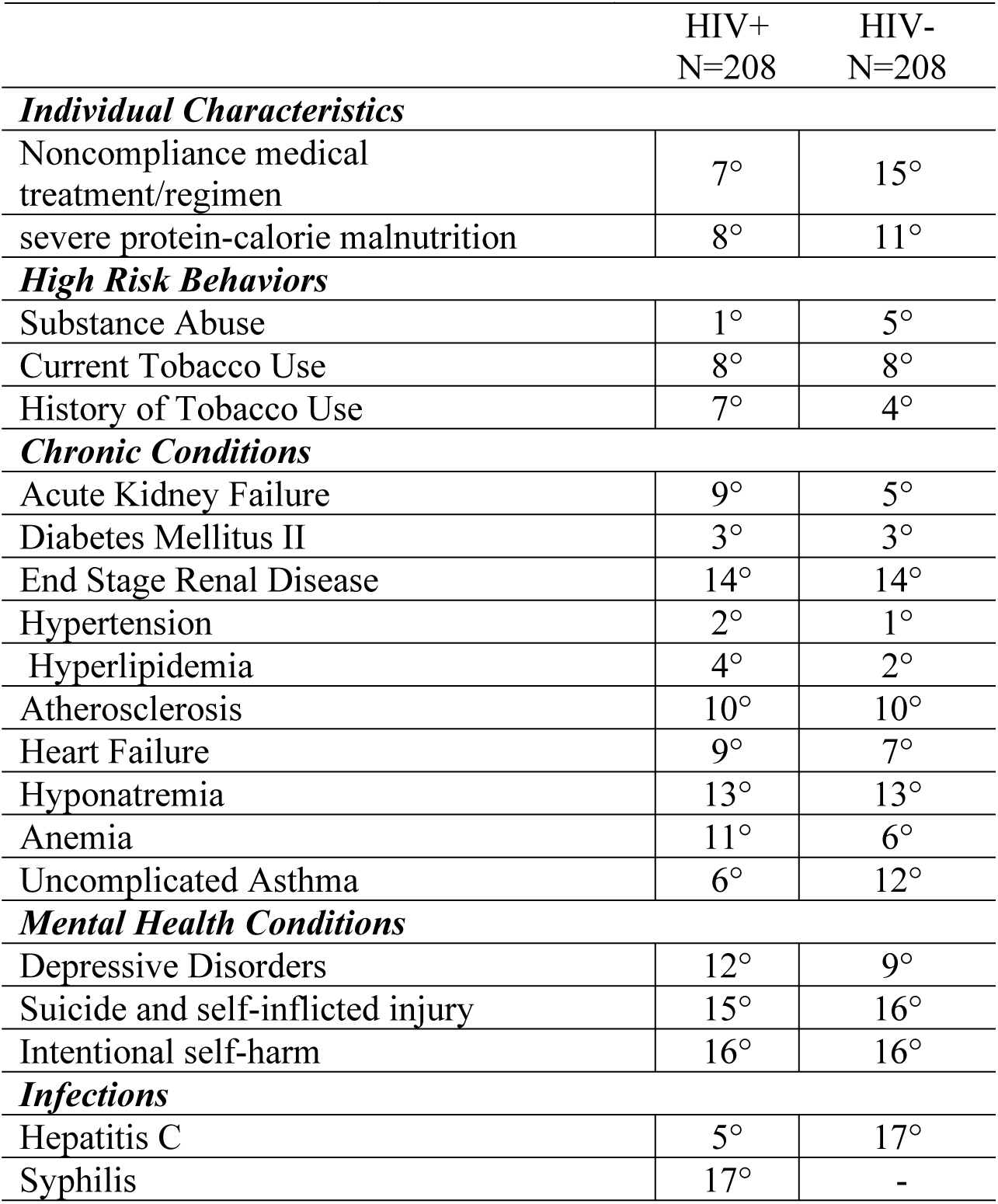
Rank (in degrees) of Factors That Contribute to Medical Resource Utilization (Charlson Scores) Based on HIV Status

**Table 2.** Descriptive Statistics, Correlation, X^2^/ t-test for Medical Resource Utilization (Charlson Scores)

|  | HIV+ | HIV- | Charlson Score |  |  |  |  |  |
| --- | --- | --- | --- | --- | --- | --- | --- | --- |
|  | (N=208) | (N=208) | Correlations |  |  | X<sup>2</sup>/ T-tests |  |  |
| Charlson Comorbidity Score | 7.42±2.4.35 | 3.52±2.97 | HIV+ |  | HIV- |  |  |  |
| **Individual Characteristics** |  |  |  |  |  |  |  |  |
| Gender |  |  |  |  |  |  |  |  |
| Male | 135 | 135 |  |  |  |  |  |  |
| Female | 73 | 73 |  |  |  |  |  |  |
| Age | 50.6±13.2 | 50.6±13.2 | 0.354 | ** | 0.515 | ** | -3.95; df=414 | ** |
| Length of Stay | 86.6±159.5 | 63.4±105 | 0.292 | ** | 0.283 | ** | -4.79; df=414<sup>a</sup> | ** |
| Admissions Frequency | 7.57±10.7 | 6.17±8.4 | 0.374 | ** | 0.248 | ** | -4.78; df=414<sup>a</sup> | ** |
| Noncompliance med treatment/regimen | 27 | 6 | 0.025 |  | -0.059 |  | 25; df=19 |  |
| severe protein-calorie malnutrition | 25 | 16 | 0.033 |  | 0.071 |  | 16.27; df=19 |  |
| **High Risk Behaviors** |  |  |  |  |  |  |  |  |
| Substance Abuse | 82 | 26 | 0.092 |  | -0.042 |  | 49.26; df=19 | ** |
| Current Tobacco Use | 25 | 20 | 0.009 |  | 0.003 |  | 24.84; df=19 |  |
| History of Tobacco Use | 27 | 38 | 0.088 |  | 0.005 |  | 24.8; df=19 |  |
| **Chronic Conditions** |  |  |  |  |  |  |  |  |
| Acute Kidney Failure | 24 | 26 | 0.384 | ** | 0.386 | ** | 88.98; df=19 | ** |
| Diabetes Mellitus II | 37 | 44 | 0.167 | * | 0.295 | ** | 55.3; df=19 | ** |
| End Stage Renal Disease | 10 | 10 | 0.139 | * | 0.143 | * | 28.31; df=19 | ** |
| Hypertension | 49 | 91 | -0.054 |  | 0.045 |  | 46.55; df=19 | * |
| Hyperlipidemia | 34 | 61 | -0.004 |  | 0.162 | * | 41.2; df=19 | ** |
| Atherosclerosis | 22 | 18 | 0.104 |  | 0.073 |  | 25.92; df=19 |  |
| Heart Failure | 24 | 22 | 0.048 |  | 0.262 | ** | 32.73; df=19 | * |
| Hyponatremia | 11 | 12 | 0.071 |  | 0.082 |  | 45.14; df=19 | ** |
| Anemia | 19 | 25 | 0.046 |  | 0.01 |  | 30.54; df=19 | * |
| Uncomplicated Asthma | 28 | 15 | -0.064 |  | -0.036 |  | 15.37; df=19 |  |
| **Mental Health Conditions** |  |  |  |  |  |  |  |  |
| Depressive Disorders | 18 | 19 | 0.061 |  | -0.157 | * | 20.28; df=19 |  |
| Suicide and self-inflicted injury | 5 | 5 | 0.079 |  | 0.025 |  | 19.88; df=19 |  |
| Intentional self-harm | 3 | 5 | -0.03 |  | -0.006 |  | 7; df=19 |  |
| **Infections** |  |  |  |  |  |  |  |  |
| Hepatitis C | 32 | 2 | 0.033 |  |  |  | 30; df=19 | * |
| Syphilis | 2 | 0 | 0.047 |  |  |  | 40.86; df=19 | ** |

<sup>a</sup> Equal variances not assumed ; \*\*p < .01, \*p < .05 for t-tests and chi square tests

Significant differences were observed from bivariate analyses between HIV infected and uninfected patients based on Charlson scores. These include the Individual Characteristics of age (t=-3.95; *df*=414), length of stay (t=-4.79; *df*=414), and admission frequency (t=- 4.78; *df*=414). They also include the High Risk Behavior of substance abuse (X^2^= 49.26; *df*=19); the Chronic Conditions of acute kidney disease (X^2^= 88.98; *df*=19), diabetes mellitus II (X^2^= 55.3; *df*=19) end stage renal disease (X^2^= 28.31; *df*=19) hypertension (X^2^= 46.55; *df*=19), hyperlipidemia (X^2^= 41.20; *df*=19), heart failure (X^2^= 32.73; *df*=19), hyponatremia (X^2^= 45.14; *df*=19) and anemia (X^2^= 30.54; *df*=19). Significant differences also included the Infections of Hepatitis C (X^2^= 30; *df*=19) and syphilis (X^2^= 40.86; *df*=19), Table 2.

Significant differences were also observed for bivariate analyses among HIV infected and uninfected patients based on age groups: under 50 years of age (<50) and 50 years of age and older (≥50). For both patient populations, these include the Health Risk Behavior of Substance Abuse: HIV (X^2^=6.8; *df*=1) uninfected (X^2^= 5.65; *df*=1) and the Chronic Conditions of Hypertension: HIV (X^2^=9.66; *df*=1) uninfected (X^2^= 9.28; *df*=1), Hyperlipidemia: HIV (X^2^=11.25; *df*=1) uninfected (X^2^= 15.17; *df*=1) and Atherosclerosis: HIV (X^2^=6.35; *df*=1) uninfected (X^2^= 3.71; *df*=1). Additionally, HIV patients included the Mental Health Condition of Intentional self-harm (X^2^=3.91; *df*=1) and the Infection of Hepatitis C (X^2^=3.75; *df*=1). Significant differences in Charlson scores were observed for uninfected patients (X^2^=7.28; *df*=1) in addition to the Chronic Condition of Uncomplicated Asthma (X^2^=5.75; *df*=1) and the Mental Health Condition of Depression (X^2^=7.61; *df*=1), Table 3.

**Table 3.**
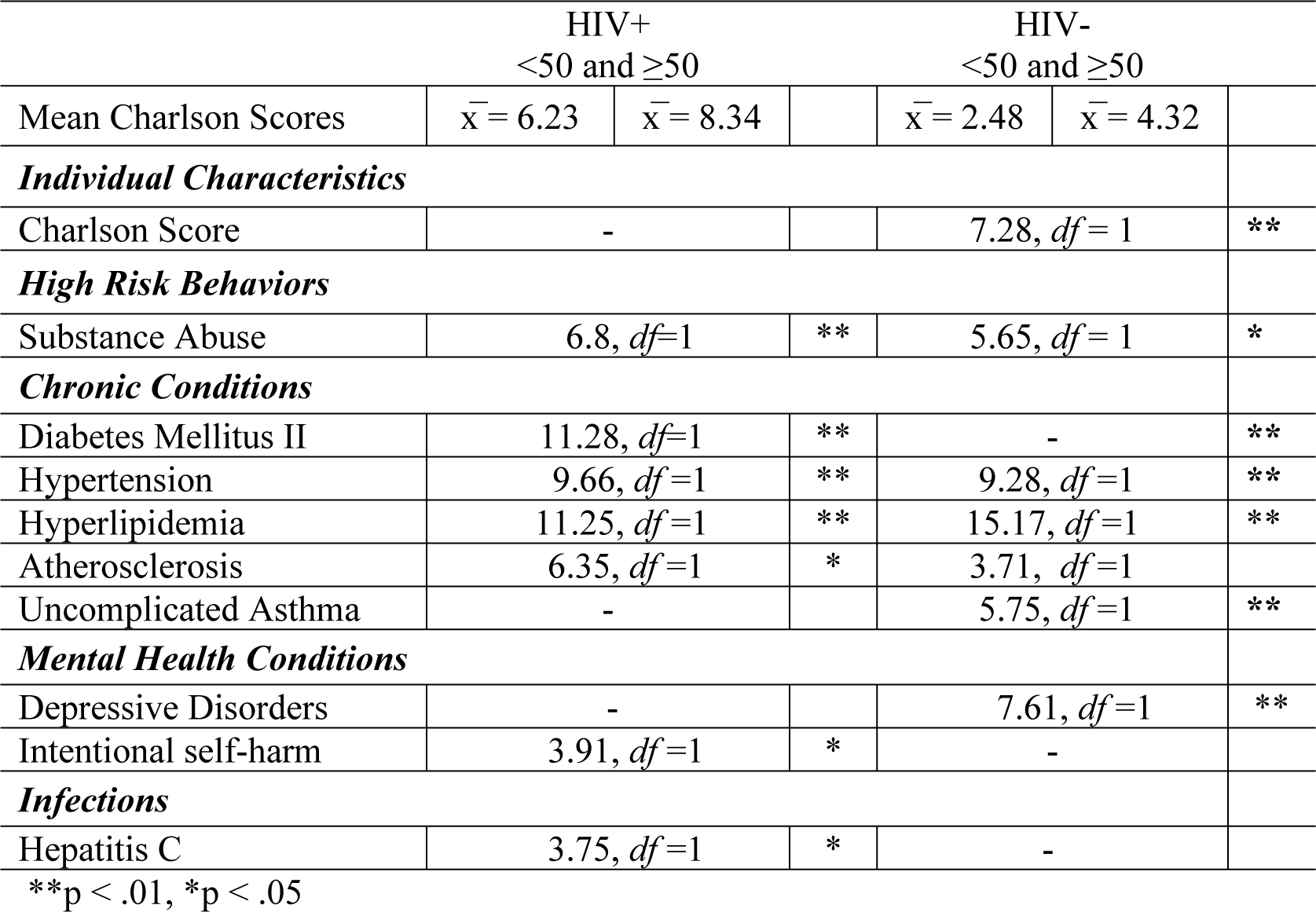
Age Differences of Factors that Contribute to Medical Resource Utilization (Charlson Scores)

The stepwise multiple regression for the HIV uninfected patients identified the individual characteristics of age (β=0.110; [95% CI 0.087, 0.133]) and length of hospital stay (β=0.006; [95% CI 0.003, 0.009]) in addition to the Chronic Conditions of acute kidney failure (β=1.556; [95% CI 0.611, 2.50]), heart failure (β= 1.713; [95% CI 0.717, 2.71]), and diabetes mellitus II (β= 1.385; [95% CI 0.634, 2.14]), as the most important contributors (p<0.05) associated with high Charlson scores, Table 4.

**Table 4.** Linear Regression Model of Best Fit for Medical Resource Utilization (Charlson Scores) (N=416)

| HIV+ Model (N=208) |  |  |  |
| --- | --- | --- | --- |
|  | Unstandardized Coefficients |  | 95.0% Confidence Interval for B |
|  | Beta |  |  |
| ***Individual Characteristics*** |  |  |  |
| Age | 0.116 | ** | 0.077 – 0.155 |
| Admission Frequency | 0.159 | ** | 0.114 – 0.205 |
| ***Chronic Conditions*** |  |  |  |
| Acute Kidney Failure | 3.27 | ** | 1.76 – 4.78 |
| Hypertension | -1.77 | * | -2.99, -0.551 |
| Diabetes Mellitus II | 1.48 | * | 0.107 – 2.85 |
| HIV- Model (N=208) |  |  |  |
| ***Individual Characteristics*** |  |  |  |
| Age | 0.110 | ** | 0.087 - 0.133 |
| Length of Stay | 0.006 | ** | 0.003 - 0.009 |
| ***Chronic Conditions*** |  |  |  |
| Acute Kidney Failure | 1.556 | ** | 0.611 - 2.50 |
| Heart Failure | 1.713 | * | 0.717 - 2.71 |
| Diabetes Mellitus II | 1.385 | * | 0.634 - 2.14 |

\*\* $p < .01$ , \* $p < .05$

The stepwise multiple regression for the HIV patients identified the individual characteristics of age (β=0.116; [95% CI 0.077, 0.155]) and admission frequency (β=0.159; [95% CI 0.114, 0.205]) in addition to the chronic conditions of acute kidney failure (β=3.27; [95% CI 1.76, 4.78]), hypertension (β= −1.77; [95% CI −2.99, −0.551]), and diabetes mellitus II (β= 1.48; [95% CI 0.107, 2.85]), as the most important contributors associated with (p<0.05) high Charlson scores, Table 4.

## Discussion

The National HIV/AIDS Strategy identified the pressing need to facilitate successful aging of PLWH [5]. As those with long-standing HIV infection age, comorbidities are becoming increasingly common [16, 24]. To support successful aging, it is essential to improve our understanding of the contribution of HIV infection to the presence of comorbidities and associated clinical outcomes. To explore this further, we utilized a clinical dataset of HIV infected and uninfected patients, matched on age and gender. We sought to identify significant population-specific differences and their contributions to medical resource utilization, informed by Charlson scores. Charlson comorbidity scores are robust predictors of mortality and medical resource utilization, [21] and an important confounding factor, essential to effective epidemiological investigations of aging and survival [6, 26]. Yet, studies utilizing Charlson scores as indicators of medical resource utilization in populations of PLWH, are sparse. Our results identified significant differences between our HIV infected and uninfected patients. Our predictive models allowed us to observe the interplay of Individual Characteristics, High Risk Behaviors, Chronic Conditions, Mental Health Conditions and Infections by HIV status.

Our regression models identified age as a high-level significant contributor to high Charlson scores in both patient populations. In HIV uninfected patients, age was the most important contributor, accounting for 50% of the model. In the HIV population, age is the second highest level contributor accounting for almost 36% of the HIV model. This indicates that unlike normal aging, other factors increase the utilization of medical resources for patients with HIV. Although our populations were matched on age, the average Charlson score was three points higher for HIV, reinforcing that the additional contributors are potential indicators of accelerated aging. Additionally, our bivariate analyses of age differences identified significant within group differences in Charlson scores for the uninfected patients, but not for the HIV patients. Similar aging-related medical resource utilization was observed in younger and older PLWH, potential evidence of accelerated aging.

Admissions frequency in HIV patients was a top predictor of high Charlson score in our regression model. In populations of PLWH, admissions type, whether urgent or non-urgent, the number of secondary diagnoses, and number of procedures, are significant contributors of medical resource utilization and significant in our model [26]. The most severe clinical conditions are often categorized by multiple diagnoses and procedures, contributing directly to length of stay; seen as a top contributor to high Charlson scores in our uninfected model [26]. Understandably, the cost of hospitalizations is a major financial burden on the US healthcare system; with HIV-related hospitalizations including the more expensive diagnostic categories [26]. Although length of stay was not significant in our HIV model, our results revealed HIV patients’ average length of stay was 87 patient days, which are 23 days more than uninfected patients. Studies have also observed that in populations of PLWH with no comorbidities, a 60% increase in length of stay and 70% increase in medical resource utilization remain, compared to uninfected groups [26].

Acute kidney failure in HIV infected and uninfected patient models as well as end stage renal disease in the HIV model were top predictors of high Charlson scores. HIV nephropathy has decreased with the use of anti-retroviral therapy. Yet, in populations of PLWH, compared to uninfected peer groups, risk factors for kidney disease were present in our models including diabetes for HIV uninfected patients and hypertension for HIV patients [23]. Other studies have also revealed that kidney-related comorbidities are associated with extremely high medical-related costs, as indicated in both our models as top contributors to high Charlson scores [27].

HIV patients are recognized to be at high risk for cardiovascular-related illness, with heart disease being a very common complication [23]. Interestingly, both models included cardiovascular related illnesses: hypertension (HIV+) and heart failure (HIV-) were top significant contributors in both groups [23]. Higher rates of hypertension are observed in PLWH in addition to associated illnesses [28, 29]. PLWH also have increased risk for cardiovascular-related mortality compared to uninfected groups. In fact, a study showed the risk of cardiovascular-related mortality increased steadily from 1999 to 2013 as the risk decreased among HIV uninfected people. Heart failure risk is also observed among PLWH with depression and hypertension [30]. Although not significant in our model, more heart failure existed in the HIV patients with hypertension, yet not the case for depression. Heart failure is also related to Diabetes Mellitus II and both diseases were present only in the HIV uninfected model. Understanding that some HIV medications increase the risk of type II Diabetes in PLWH in addition to the regular risk factors, it was surprising it did not contribute to high medical resource utilization as with our uninfected population [31, 32].

## Limitations

Our cross-sectional study explored the top diagnoses and procedure codes. Future studies on predictive modeling should explore additional clinical factors and apply other analytic approaches to support a comprehensive evaluation of medical resource utilization for effective integrated health services delivery. We did not match our sample on other factors including socioeconomic status, due to incomplete sociodemographics in the patient records. Data that comprise EHRs is collected during clinical care and not collected for research purposes. Therefore, incomplete sociodemographics will exist for a variety of indicators including race. Our cross-sectional analysis was unable to include HIV-related contributions (e.g., immune status) to the development or severity of comorbidities, only their documented presence or absence. Future longitudinal studies should account for these additional factors, and track such factors (e.g., immune status) over time to explore their contributions to comorbidity development and medical resource utilization.

Our analysis also did not include an exploration of the contribution of HIV-related clinical indicators (CD4, viral load, antiretrovirals); the analysis of HIV-related clinical indicators one point in time (cross-sectional analysis) would not be informative to the presence and absence of the observed comorbidities. Future longitudinal studies should analyze HIV-related clinical indicators over time, as medications, including protease inhibitors, are linked to increase risk of cardiovascular related illness. This should also be explored in future predictive modeling studies [23].

## Conclusions

As HIV population’s age both locally and globally, preventing, identifying and managing comorbidities is increasingly important. An essential need exists to further understand the causal factors of identified aging outcomes and in the exploration of additional clinical indicators to inform optimal treatment, care and self-management. Interventions targeting aging phenotypes, specific to HIV, have been sparse. Additional contributions to the development of such phenotypes, were found in our study and critical as PLWH age. Similarities and differences were observed between our age and gender matched patients and factor-specific contributions in higher medical resource utilization observed. Our findings add to the literature on HIV and aging-related outcomes and support HIV and aging phenotype development.

## List of Abbreviations

PLWH: People Living with HIV/AIDS
PPMC: Pearson Product Moment Correlations
CI: Confidence Intervals

## Acknowledgements

This work was supported by Columbia University’s Office of the Vice Provost for Diversity and Inclusion

## Authors Contributions

Authors assisted in the data interpretation (MO, SY), manuscript drafting (MO), revising intellectual content (MO, SY). This manuscript has not been published and is not under consideration for publication elsewhere.

